# Serum Metabolites Associated with Brain Amyloid Beta Deposition, Cognitive Dysfunction, and Alzheimer’s Disease Progression

**DOI:** 10.1101/2020.11.25.394262

**Authors:** Kwangsik Nho, Alexandra Kueider-Paisley, Matthias Arnold, Siamak MahmoudianDehkordi, Shannon L. Risacher, Gregory Louie, Colette Blach, Rebecca Baillie, Xianlin Han, Gabi Kastenmüeller, P. Murali Doraiswamy, Rima Kaddurah-Daouk, Andrew J. Saykin, for the Alzheimer’s Disease Neuroimaging Initiative and the Alzheimer Disease Metabolomics Consortium

## Abstract

**RATIONALE:** Metabolomics in the Alzheimer’s Disease Neuroimaging Initiative (ADNI) cohort provides a powerful tool for mapping biochemical changes in AD, and a unique opportunity to learn about the association between circulating blood metabolites and brain amyloid-β deposition in AD.

**OBJECTIVES:** We examined 140 serum metabolites and their associations with brain amyloid-β deposition, cognition, and conversion from mild cognitive impairment (MCI) to AD.

**FINDINGS:** Serum-based targeted metabolite levels were measured in 1,531 ADNI participants. We performed association analysis of metabolites with brain amyloid-β deposition measured from [18F] Florbetapir PET scans. We identified nine metabolites as significantly associated with amyloid-β deposition after FDR-based multiple comparison correction. Higher levels of one acylcarnitine (C3; propionylcarnitine) and one biogenic amine (kynurenine) were associated with decreased amyloid-β accumulation. However, higher levels of seven phosphatidylcholines (PC) were associated with increased amyloid deposition. In addition, PC ae C44:4 was significantly associated with cognition and conversion from MCI to AD dementia.

**CONCLUSION:** Perturbations in PC and acylcarnitine metabolism may play a role in features intrinsic to AD including amyloid-β deposition and cognitive performance.

## Introduction

Late-onset Alzheimer’s disease (AD) is an age-related neurodegenerative disease with a long preclinical period extending at least two decades (1). Understanding the etiology of AD has proved to be challenging due to the complexity of the disease including its long pre-symptomatic period and variability in clinical symptoms. Targeted and non-targeted metabolomic and lipidomic platforms are being used to investigate the molecular underpinnings of pathological hallmarks of AD relevant to the pathophysiology of the disease in greater detail. Compared to cognitively normal controls, AD patients have impairments in phospholipid homeostasis (2–5), up-regulated degradation of membrane phospholipids and sphingolipids (2, 6–8), and impairments in neurotransmission (6, 7). Further, several metabolomics studies report overlap between associations of multiple metabolite classes (e.g., phospholipids, sphingomyelins, acylcarnitines, branched chain and aromatic amino acids) with known risk factors of AD and cognitive impairment including insulin resistance (9–12).

Although there is extensive literature related to metabolic perturbations in AD and its risk factors using metabolomic platforms, it is still unclear how circulating serum metabolites are associated with brain amyloid-β deposition. Evidence suggests there are metabolic interactions between peripheral and central compartments that go beyond substrate transport (5, 13–17), however, our understanding of these interactions is limited. Work by the Alzheimer’s Disease Metabolomics Consortium (ADMC) identified the role of the gut microbiome and liver in cognitive decline and changes in the brain that are hallmarks of the disease, highlighting the cross-talk between peripheral and central compartments (13–15). Amyloid deposition is one of the central neuropathological features of AD, yet little is known about its association with the metabolome. Understanding the associations between circulating metabolites and amyloid-β deposition could shed light on the mechanisms underlying the association between metabolic perturbations and AD risk as well as lead to the identification of novel biomarkers. Here, we performed association analyses of circulating serum metabolites with measures of amyloid-β deposition, cognition, and conversion to AD dementia from mild cognitive impairment (MCI) in a large national multi-center cohort, the Alzheimer’s Disease Neuroimaging Initiative (ADNI).

## Results

### Study sample

Demographic information for all diagnosis groups is presented in **Supplementary Table 1**. This included 1,531 Alzheimer’s Disease Neuroimaging Initiative (ADNI) participants (370 cognitively normal older adult controls (CN), 95 with significant memory concern (SMC), 271 with early mild cognitive impairment (EMCI), 491 with late MCI (LMCI), and 304 with AD).

### Region of Interest (ROI) based analysis of amyloid-β PET

Using 783 ADNI participants (167 CN, 75 SMC, 267 EMCI, 146 LMCI, 128 AD) with both P180 metabolites and [^18^F] Florbetapir PET scans, we evaluated whether metabolites were associated with ROI-based brain amyloid-β load by performing an association analysis with each metabolite. We used global cortical amyloid deposition measured by amyloid PET scans. We identified nine metabolites significantly associated with global cortical amyloid deposition after applying FDR-based multiple comparison correction (**Table 1**). Increased levels of one acylcarnitine (C3; propionylcarnitine) and one biogenic amine (kynurenine) were associated with decreased brain amyloid-β accumulation; whereas increased levels of seven phosphatidylcholines ((PC); lysoPC a C18:2, PC aa C42:0, PC ae C42:3, PC ae C44:3, PC ae C44:4, PC ae C44:5, and PC ae C44:6) were associated with increased brain amyloid-β accumulation.

**Table 1.** Results of association of P180 metabolites with a global cortical amyloid deposition measured from amyloid PET scans (FDR-corrected *p*-value < 0.05)

| Metabolite | $\beta$ (SE) | Corrected $p$-value |
| --- | --- | --- |
| C3 | -0.009 (0.003) | 0.042 |
| Kynurenine | -0.010 (0.003) | 0.036 |
| LysoPC a C18:2 | 0.009 (0.003) | 0.042 |
| PC aa C42:0 | 0.009 (0.003) | 0.042 |
| PC ae C42:3 | 0.009 (0.003) | 0.042 |
| PC ae C44:3 | 0.010 (0.003) | 0.036 |
| PC ae C44:4 | 0.008 (0.003) | 0.042 |
| PC ae C44:5 | 0.008 (0.003) | 0.042 |
| PC ae C44:6 | 0.009 (0.003) | 0.036 |

### Detailed whole brain analysis of amyloid-β PET

In addition to the priori region-based amyloid-β PET analysis, we performed a detailed whole-brain analysis of brain amyloid-β accumulation on a voxel-wise level measured from amyloid-β PET scans for nine metabolites (C3, kynurenine, lysoPC a C18:2, PC aa C42:0, PC ae C42:3, PC ae C44:3, PC ae C44:4, PC ae C44:5, and PC ae C44:6) that were significantly associated with a global cortical amyloid deposition. We used a general linear model approach to identify brain regions where amyloid-β accumulation was associated with levels of each metabolite. The results of the voxel-wise association between brain amyloid-β load and metabolites are shown in **Fig. 1**. Higher levels of C3 were associated with decreased amyloid-β accumulation in the bilateral frontal and temporal lobes, with a global maximum association in the left temporal cortex (Brodmann area (BA) 38) (**Fig. 1a**). Higher levels of kynurenine were associated with decreased amyloid-β accumulation in a widespread pattern, especially the bilateral temporal, parietal, and frontal lobes, with a global maximum association in the left dorsolateral prefrontal cortex (BA 9) (**Fig. 1b**). In contrast, higher levels of lysoPC a C18:2. PC aa C 42:0, PC ae C42:3, PC ae C44:3, PC ae C44:4, PC ae C44:5, and PC ae C44:6 were associated with increased amyloid-β accumulation in a widespread pattern of significant voxels, especially in the bilateral frontal, temporal, and parietal lobes. Global maximum associations were noted in the right parietal cortex (BA 7) (**Fig. 1c**) for lysoPC a C18:2, the right dorsolateral prefrontal cortex (BA 9) (**Fig. 1d**) for PC aa C42:0, the right parietal cortex (BA 39) (**Fig. 1e**) for PC ae C42:3, the right primary auditory cortex (BA 41) (**Fig. 1f, 1h, 1i**) for PC ae C44:3 and PC ae C44:5, and the left parietal cortex (BA 40) (**Fig. 1g**) for PC ae C44:4.

**Figure 1.**
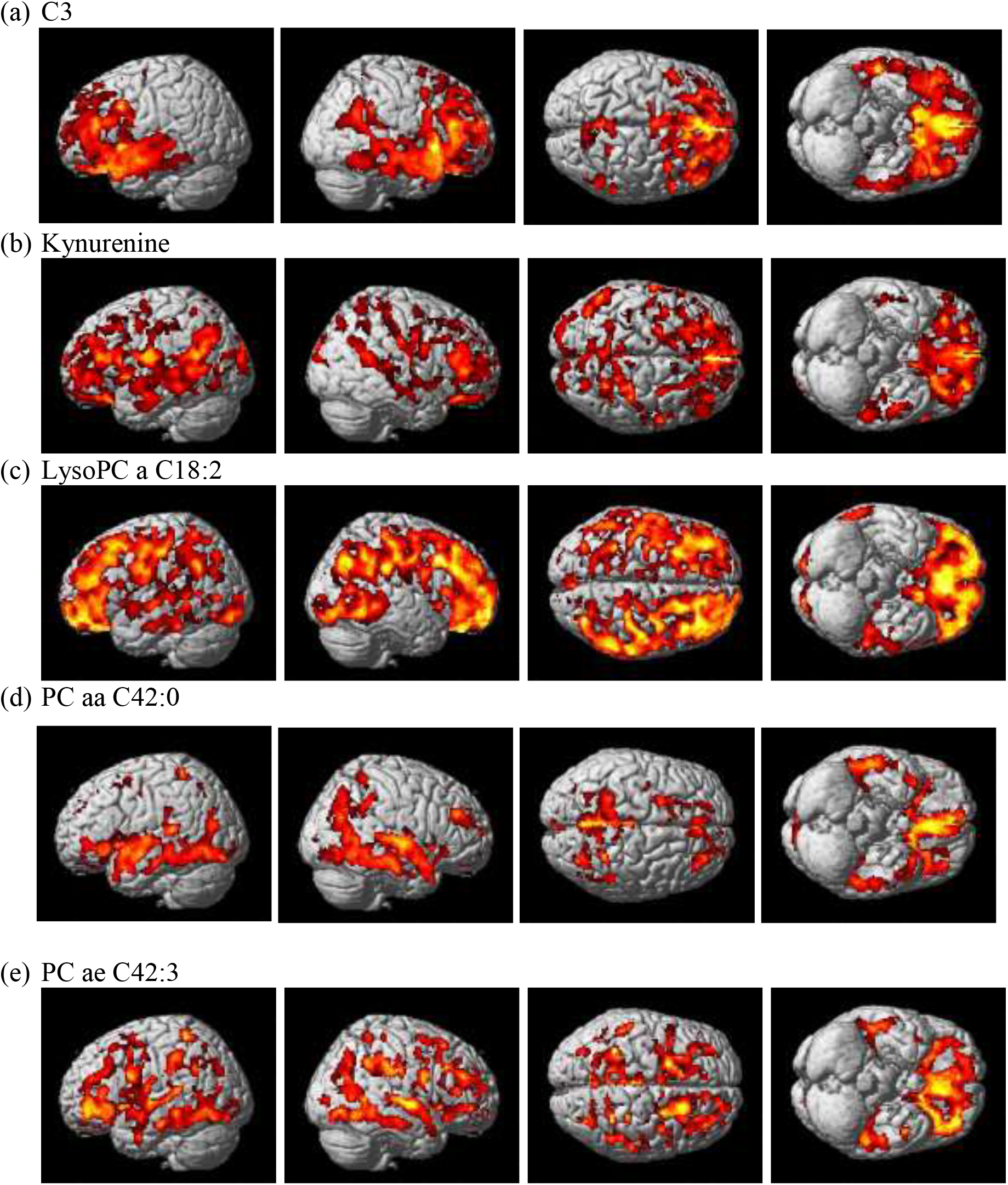
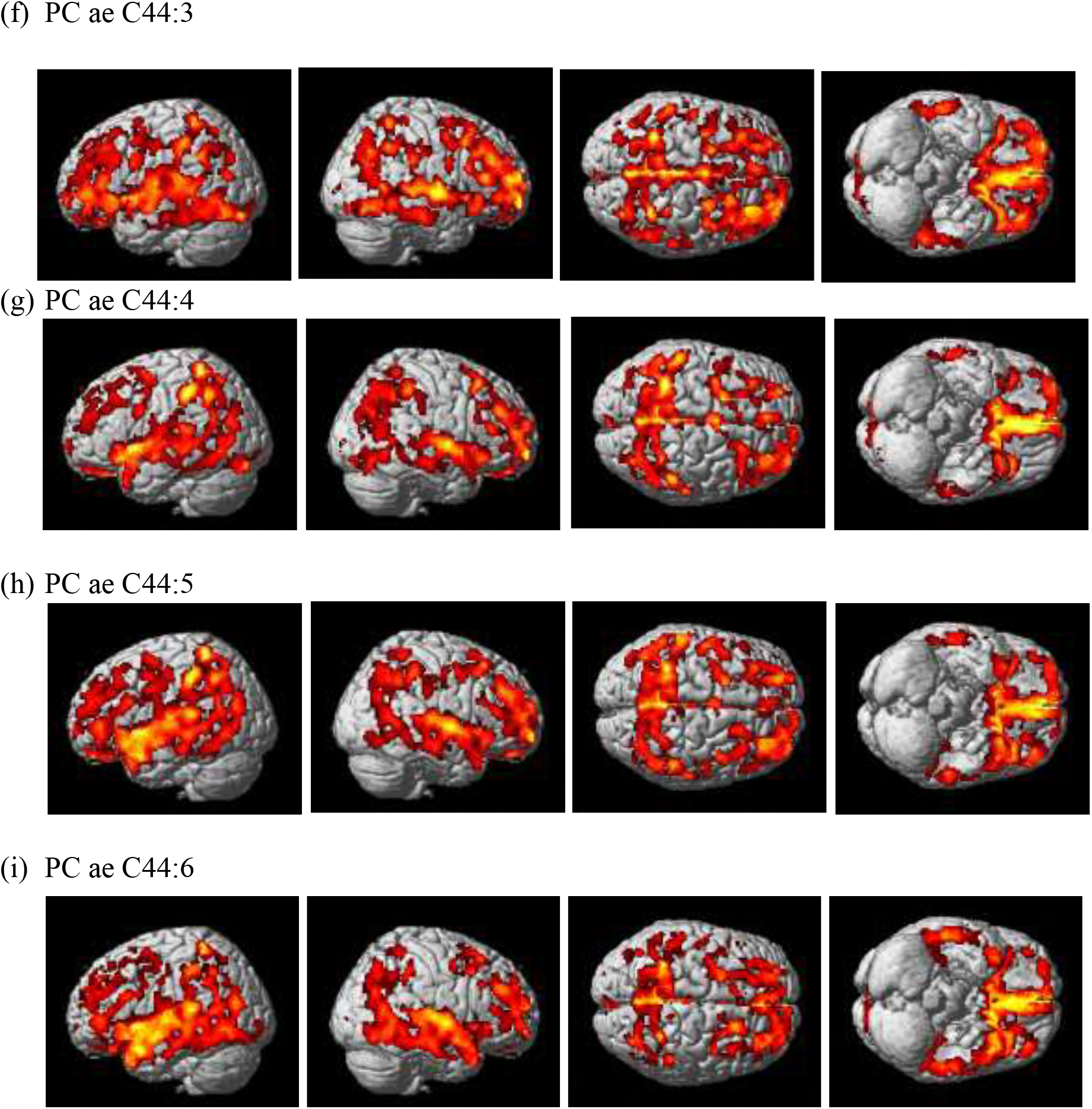
Detailed whole-brain voxel-based imaging analysis for amyloid-β accumulation using [^18^F]Florbetapir PET scans (corrected *p*-value < 0.05). Higher levels of (a) C3 and (b) kynurenine were associated with decreased amyloid-β accumulation. Higher levels of (c) lysoPC a C18:2, (d) PC aa C42:0, (e) PC ae C42:3, (f) PC ae C44:3, (g) PC ae C44:4, (h) PC ae C44:5, and (i) PC ae C44:6 were associated with increased amyloid-β accumulation.

### Association of significant metabolites with cognition

Using composite scores for memory (ADNI-MEM) and executive functioning (ADNI-EF), we performed a further analysis of nine metabolites (C3, kynurenine, lysoPC a C18:2, PC aa C42:0, PC ae C42:3, PC ae C44:3, PC ae C44:4, PC ae C44:5, and PC ae C44:6) that were significantly associated with a global cortical amyloid deposition. We identified significant associations of the selected metabolites with cognition after adjusting for multiple comparison correction using FDR (**Table 2**). Higher levels of C3 and kynurenine were associated with higher memory scores. In contrast, higher levels of PC ae C44:4 and PC ae C44:6 were associated with lower memory scores. Higher kynurenine levels were associated with higher executive functioning scores, whereas higher levels of PC aa C42:0, PC ae C44:4, PC ae C44:5, and PC ae C44:6 were associated with lower executive functioning scores.

**Table 2.** Results of association of the nine metabolites significantly associated with brain amyloid-β deposition with composite cognitive performance measures

| Metabolite | Memory Composite Score |  | Executive Function Composite Score |  |
| --- | --- | --- | --- | --- |
| | $\beta$ (SE) | Corrected $p$-value | $\beta$ (SE) | Corrected $p$-value |
| C3 | 0.07 (0.02) | 0.016 | 0.05(0.03) | 0.148 |
| Kynurenine | 0.05 (0.02) | 0.042 | 0.07 (0.03) | 0.021 |
| LysoPC a C18:2 | -0.01 (0.02) | 0.893 | 0.04 (0.03) | 0.213 |
| PC aa C42:0 | -0.05 (0.02) | 0.074 | -0.08 (0.03) | 0.016 |
| PC ae C42:3 | -0.0004 (0.02) | 0.987 | -0.004 (0.03) | 0.940 |
| PC ae C44:3 | -0.03 (0.02) | 0.263 | -0.05 (0.03) | 0.132 |
| PC ae C44:4 | -0.06 (0.02) | 0.038 | -0.08 (0.03) | 0.015 |
| PC ae C44:5 | -0.03 (0.02) | 0.226 | -0.08 (0.03) | 0.015 |
| PC ae C44:6 | -0.05 (0.02) | 0.040 | -0.10 (0.03) | 0.004 |

### Association of significant metabolites with conversion to AD dementia in MCI

Of the 615 MCI patients with both P180 metabolites and a follow-up clinical diagnosis, 182 MCI patients progressed to AD dementia over the two year period from the baseline visit (MCI-Converter) and 433 MCI patients remained stable over two year period from the baseline visit (MCI-Stable). We performed an association of levels of the nine identified metabolites at baseline with MCI-to-AD dementia conversion over two years (MCI-Stable and MCI-Converter). Only PC ae C44:4 showed a significant group difference between MCI-Stable and MCI-Converter groups after FDR-correction (**Fig. 2**). The MCI-Converter group has significantly higher levels of PC ae C44:4 at baseline compared to the MCI-Stable group.

**Figure 2.**
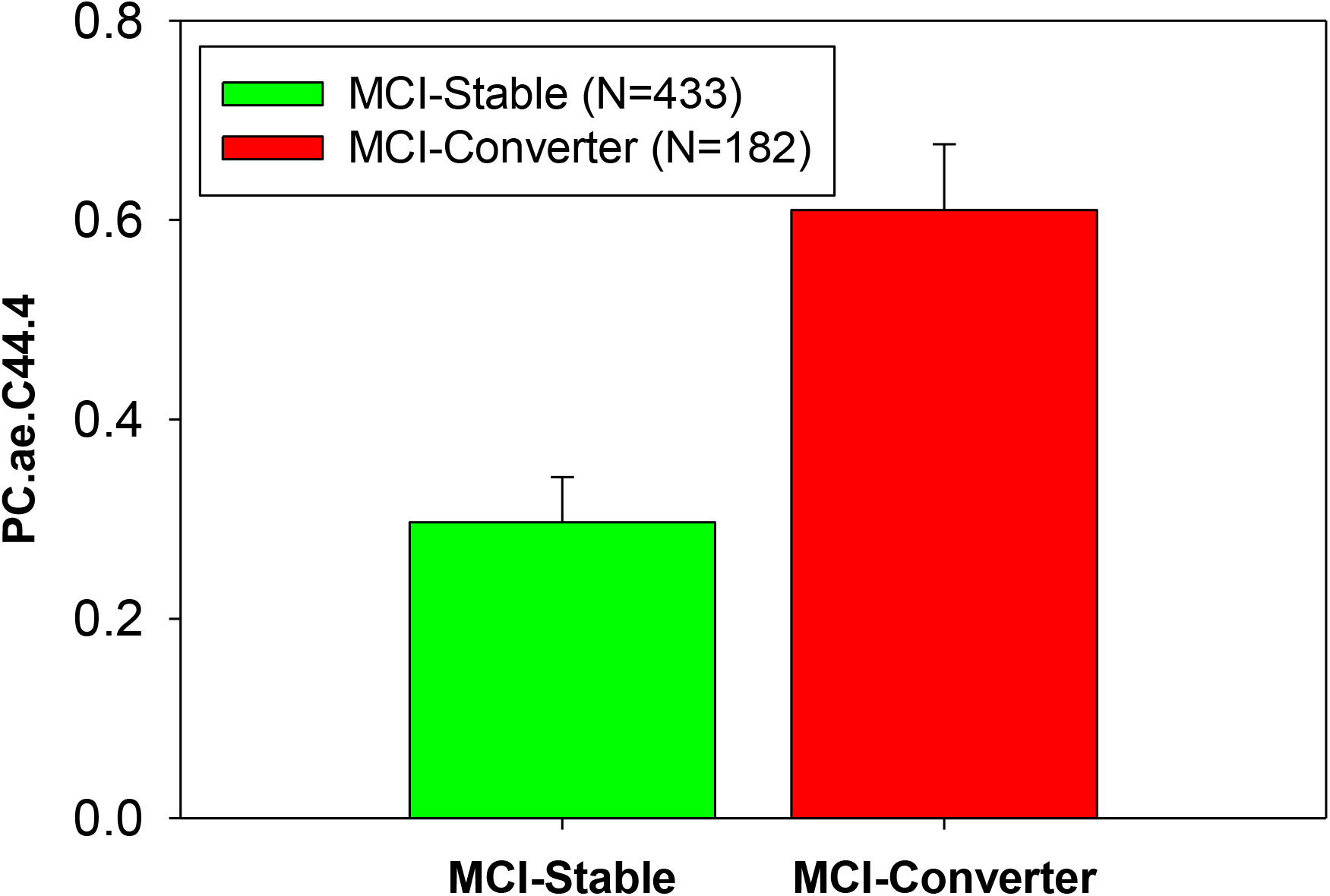
Group comparison of Mild Cognitive Impairment groups (MCI-Stable and MCI-Converter) based on baseline diagnosis and two year conversion status for PC ae C44:4.

## Discussion

In this report we analyzed 140 targeted serum-based metabolites in the ADNI cohort to investigate the relationship between metabolite levels and amyloid-β deposition, cognition, and disease progression. We identified nine metabolites (C3, kynurenine, lysoPC a C18:2, PC aa C42:0, PC ae C42:3, PC ae C44:3, PC ae C44:4, PC ae C44:5, and PC ae C44:6) significantly associated with global cortical amyloid deposition including glycerophospholipids, an acylcarnitine, and a biogenic amine. Of the nine amyloid-β-associated metabolites, six were significantly associated with cognitive performance (C3, kynurenine, PC aa C42:0, PC ae C44:4, PC ae C44:5, and PC ae C44:6), and one metabolite (PC ae C44:4) was associated with conversion to AD dementia from MCI.

### Phosphatidylcholines

Our findings suggest that dysregulation of peripheral PC metabolism is associated with earlier pathological changes noted in AD as measured by amyloid-β deposition as well as later clinical changes including changes in memory and executive functioning. Higher levels of seven PCs were associated with increased amyloid deposition (lysoPC a C18:2, PC aa C42:0, PC ae C42:3, PC ae C44:3, PC ae C44:4, PC ae C44:5, and PC ae C44:6). Of these seven PCs, four were significantly associated with cognitive performance. Higher levels of PC ae C44:4 and PC ae C44:6 were associated with lower memory scores, whereas higher levels of PC aa C42:0, PC ae C44:4, PC ae C44:5, and PC ae C44:6 were associated with lower executive functioning scores. The MCI-Converter group had significantly higher levels of PC ae C44:4 at baseline compared to the MCI-Stable group.

In our previous publication using p180 data from the ADNI cohort, we reported PC ae C44:4, PC ae C44:5, and PC ae C44:6 were all associated with CSF Aβ pathology, and PC ae C44:4 and PC ae C44:5 were further associated with brain glucose metabolism as measured by FDG PET (18). Effect directions were consistent with our current findings. Other studies reported associations of PCs with changes in cognition that become apparent later in the disease (19–21). In addition, there is research to suggest PC metabolism is perturbed in AD. Gonzales et al. reported an increase of PCs in serum samples of patients with AD (2). Results from previous studies suggest dysregulation in the biosynthesis, turnover, and acyl chain remodeling of phospholipids that is in line with increased phospholipid breakdown due to phospholipase A2 (PLA2) over activation (2, 22, 23). Perturbations in PC metabolism appear to play a role in several key molecular pathways intrinsic to cognitive decline and AD including neuroinflammation through arachidonic acid signaling, amyloid precursor protein processing through phospholipase A2, and cholesterol transport through high-density lipoproteins (24–26). Further, PCs play a critical role in the balance between cell proliferation and death which has clear implications for the pathogenesis of AD (27). Ether-linked PCs may be located in membrane rafts and could support the hypothesis that lipid rafts play a critical role in AD through the promotion of amyloid-β peptide and its aggregation (28). Amyloid-β peptides are derived from the proteolytic processing of amyloid precursor protein within lipid rafts (29). Abnormal lipid rafts have been noted in post-mortem brain samples from AD patients (30). While these associations with lipid metabolites and amyloid-β accumulation may shed led of mechanisms related to Aβ pathology in AD, given the limitations of the current study including its cross-sectional design and measurement of blood in the periphery, we cannot make assumptions about directionality.

LysoPC C18:2 was significantly associated with increased amyloid accumulation in a widespread pattern especially in the bilateral frontal, temporal, and parietal lobes. LysoPCs are known to play a role in inflammation (31) and have both pro- and anti-atherogenic properties (32). Low levels of plasma lysoPC 18:2 predict impaired glucose tolerance, insulin resistance (33), type 2 diabetes (34), coronary artery disease, and memory impairment (35) as well as declines in gait speed in older adults (36), all of which are known to be associated with AD. Amyloid-β may directly disrupt integrity of the lipid bilayer by interactions with phospholipids (37).

### Propionylcarnitine

We found that higher levels of propionylcarnitine (C3) were associated with less amyloid-β deposition in the bilateral temporal and frontal lobes. Previous studies reported perturbations in acylcarnitines in AD and MCI patients compared to cognitively normal controls as well as diagnostic converters(18, 21, 35, 38). In our previous publication examining effects of sex and APOE ɛ4 on the metabolome using ADNI data (18) we reported significant sex differences in MCI patients in multiple acylcarnitines (i.e., C0, C3, C9, and C18:2). Mapstone and colleagues reported significantly lower plasma levels of C3 in the converter group, before and after phenoconversion to MCI/AD, and in the MCI/AD group (35). In a separate study, levels of C3 were significantly lower in MCI compared to cognitively normal controls (38).

Acylcarnitines are involved in energy metabolism, mitochondrial function, neurotransmission, and neuroprotection (39). Acylcarnitines directly reflect the oxidation rate of amino acids and fatty acids (40–43). González-Dominguez et al. (44) demonstrated the deficit of several acylcarnitines, including C3, in the brain (hippocampus, cortex, and cerebellum) of APP/PS1 transgenic mouse model of AD. Short- and medium-chain acyl groups are catalyzed by acyltransferases that are located in microsomes and peroxisomes whereas long-chain acyl groups are catalyzed by carnitine palmitoyltransferase I and II located on the mitochondria membranes.

Our results could suggest selective perturbed metabolism of short-chain acylcarnitines in the peroxisomes (43, 45). Propionylcarnitine reflects the propionyl CoA pool which is a byproduct of isoleucine and valine catabolism. A direct link between branch chain amino acids (BCAA) and C3 has been demonstrated by a rise in circulating levels of C3 in response to BCAA supplementation (46). We and others have implicated BCAA in cognitive decline, AD diagnosis, and AD risk (18, 21, 47–49)

### Kynurenine

We report that higher levels of kynurenine were associated with widespread decreased amyloid-β accumulation, especially in the bilateral temporal, parietal, and frontal lobes as well as higher memory and executive function scores. An independent study of older adults with normal global cognition reported a positive correlation between plasma Aβ_1-42_ and metabolites in the kynurenine pathway(50). These same metabolites were also associated with plasma neurofilament light chain (NFL), an emerging marker of neurodegeneration and inflammation. (50). Plasma NFL, a marker of axonal cytoskeletal damage, and plasma Aβ have been shown to reflect brain Aβ deposition (51, 52). However, it is possible that axonal damage may be due to other etiologies related to axonal injury including vascular pathologies or traumatic brain injuries. The kynurenine pathway is dysregulated in neuroinflammation, and chronic activation of the kynurenine pathway has been implicated in AD as well as other neuropsychiatric disorders including depression, suicidality, and schizophrenia (53–58). Neuropsychiatric symptoms are common among AD patients with up to 80% reporting at least one neuropsychiatric symptom (59). This could suggest that in AD, the neuroinflammatory consequences of chronic activation of the kynurenine pathway could be a factor in cognitive changes and/or neuropsychiatric symptoms noted in the disease. Metabolites in the kynurenine pathway, including kynurenine, readily cross the blood brain barrier suggesting that circulating levels may contribute significantly to cerebral pools (60). However, it should be noted that while mechanistic data (e.g., quinolinic acid neurotoxicity and 3-hydroxyanthranilic acid mediated reactive oxidative species production (61, 62)) support the role of kynurenine pathway metabolites in AD pathogenesis, it is still unclear if these are causative of disease or merely the result of chronic neuroinflammation.

## Limitations

As metabolomics data are only available at one time point, current analyses are cross-sectional and therefore we were unable to explore issues related to causality. Due to the sample recruitment strategy used in the ADNI cohort, which was designed to model a simulated clinical trial for MCI and AD, participants were selected for symptoms and early stage pathology and are likely to differ in significant ways from those recruited into population-based cohorts. As data from both designs become available, the difference between these two sampling strategies is likely to reveal important factors related to early disease and the role of comorbidities.

## Conclusions

The present study provides insight into the association of serum-based circulating metabolites with brain amyloid-β accumulation across the whole brain. Perturbations in phosphatidylcholine metabolism may point to issues with membrane restructuring leading to the accumulation of amyloid-β in the brain. Additional studies are needed to explore whether these metabolites play a causal role in the pathogenesis of Alzheimer’s disease or if they are biomarkers for systemic changes during preclinical phases of the disease.

## Methods

### Study cohort

Serum samples and data analyzed in the present report were obtained from the Alzheimer’s Disease Neuroimaging Initiative (ADNI) cohort. The initial phase (ADNI-1) was launched in 2003 to test whether serial magnetic resonance imaging (MRI), position emission tomography (PET), other biological markers, and clinical and neuropsychological assessment could be combined to measure the progression of mild cognitive impairment (MCI) and early AD. ADNI-1 was extended to subsequent phases (ADNI-GO, ADNI-2, and ADNI-3) for follow-up for existing participants and additional new enrollments. Inclusion and exclusion criteria, clinical and neuroimaging protocols, and other information about ADNI can be found at www.adni-info.org (63, 64) Demographic information, raw neuroimaging scan data, *APOE*, and clinical information are available and were downloaded from the ADNI data repository (www.loni.usc.edu/ADNI/). Written informed consent was obtained at the time of enrollment that included permission for analysis and data sharing and consent forms were approved by each participating sites’ Institutional Review Board (IRB).

### AbsoluteIDQ-p180 kit metabolites

Metabolites were measured using a targeted metabolomics approach using the AbsoluteIDQ-p180 kit (BIOCRATES Life Science AG, Innsbruck, Austria), with an ultra-performance liquid chromatography (UPLC)/MS/MS system [Acquity UPLC (Waters), TQ-S triple quadrupole MS/MS (Waters)], which provides measurements of up to 186 endogenous metabolites quantitatively (amino acids and biogenic amines) and semi-quantitatively (acylcarnitines, sphingomyelins, PCs, and lyso-glycero-phosphatidylcholines (a = acyl) [lysoPCs] across multiple classes). The AbsoluteIDQ-p180 kit has been fully validated according to European Medicine Agency Guidelines on bioanalytical method validation. In addition, plates include an automated technical validation to approve the validity of the run and provide verification of the actual performance of the applied quantitative procedure including instrumental analysis. The technical validation of each analyzed kit plate was performed using MetIDQ software based on results obtained and defined acceptance criteria for blank, zero samples, calibration standards and curves, low/medium/high-level QC samples, and measured signal intensity of internal standards over the plate (21).

Metabolomics data and pre-processed data are accessible through the AMP-AD Knowledge Portal (https://ampadportal.org). The AMP-AD Knowledge Portal is the distribution site for data, analysis results, analytical methodology and research tools generated by the AMP-AD Target Discovery and Preclinical Validation Consortium and multiple Consortia and research programs supported by the National Institute on Aging. Information on data availability and accessibility is available in the Data Availability section.

### P180 quality control

Metabolites with >40% of measurements below the lower limit of detection (LOD) were excluded from the analysis. Metabolite values were scaled across the different plates using the quality control (QC) pool duplicates. Metabolite measurements below LOD were imputed using each metabolite’s LOD/2 value. Using the blinded duplicates, we selected metabolites with a coefficient of variation <20% and an intraclass correlation coefficient >0.65 (21). We checked for the presence of multivariate outlier participants by evaluating the first and second principal components in each platform. For the participants with duplicated measurements, we used the average values of the two measured values in further analyses. The QC process resulted in 140 metabolites for further analysis. The QC-passed preprocessed metabolite measurements were adjusted for the effect of medication use at baseline on metabolite levels (see Toledo et al. 2017 (21) for adjustment description details).

#### Neuroimaging processing

### Positron Emission Tomography (PET)

Pre-processed [^18^F] Florbetapir PET scans (co-registered, averaged, standardized image and voxel size, uniform resolution) were downloaded from the ADNI LONI site (http://adni.loni.usc.edu) as described in previously reported methods for acquisition and processing of PET scans from the ADNI sample (65, 66). For [^18^F] Florbetapir PET, scans were intensity-normalized using a whole cerebellum reference region to create SUVR images.

#### Statistical analyses

For [^18^F] Florbetapir PET, a mean SUVR value was extracted using MarsBaR from a global cortical region generated from an independent comparison of ADNI-1 [11C]Pittsburgh Compound B SUVR scans (regions where AD > CN).

We performed a linear regression association analysis of P180 metabolites with brain amyloid-β deposition and with composite scores for memory and executive functioning. In addition, we explored the group differences between MCI-Converter and MCI-Stable groups stratified by baseline diagnosis and two year conversion from MCI to AD. Of the 615 MCI patients with both P180 metabolites and a follow-up clinical diagnosis, 182 MCI patients progressed to AD dementia over the two year period from the baseline visit (MCI-Converter) and 433 MCI patients remained stable over two year period from the baseline visit (MCI-Stable). Age, sex, years of education, body mass index (BMI), study phase (ADNI-1 or ADNI-GO/2), and *APOE* ∊4 status were used as covariates. FDR-based multiple comparison adjustment with the Benjamini-Hochberg procedure was used because the metabolites and AD biomarker phenotypes were strongly correlated with each other (67). Not accounting for this high collinearity of dependent variables would lead to an overly stringent correction for multiple testing.

### Whole brain imaging analysis

The processed [^18^F] Florbetapir PET images were used to perform a voxel-wise statistical analysis of the effect of P180 metabolite levels on amyloid-β accumulation across the whole brain using SPM8 (www.fil.ion.ucl.ac.uk/spm/). We performed a multivariable regression analysis using age, sex, BMI, *APOE* ∊4 status, and study phase (ADNI-1 or ADNI-GO/2) as covariates. In the voxel wise whole brain analysis, the significant statistical parameters were selected to correspond to a threshold of *p* < 0.05 (FDR-corrected).

## Data Availability

Metabolomics datasets from the AbsoluteIDQ-p180 metabolomics kit used in the current analyses for the ADNI-1 and ADNI-GO/-2 cohorts are available via the Accelerating Medicines Partnership-Alzheimer’s Disease (AMP-AD) Knowledge Portal and can be accessed at http://dx.doi.org/10.7303/syn5592519 (ADNI-1) and http://dx.doi.org/10.7303/syn9705278 (ADNI-GO/-2). The full complement of clinical and demographic data for the ADNI cohorts are hosted on the LONI data sharing platform and can be requested at http://adni.loni.usc.edu/data-samples/access-data/.

## Author Contributions

Nho, Arnold had full access to all of the data in the study and take responsibility for the integrity of the data and the accuracy of the data analysis.

**Statistical analyses also included**: Arnold, Kastenmüller, MahmoudianDehkordi

**Data management**: Blach

**Concept and design**: Kaddurah-Daouk led concept and design team that included all co-authors. She leads the ADMC that led the study and generated data. Saykin designed the strategy to map metabolites to AD biomarkers.

**Drafting of the manuscript**: Kueider-Paisley, Nho

**Biochemical interpretation**: Kaddurah-Daouk, Kastenmüller, Baillie, Han, Risacher, Saykin

**Data deposition**: Alzheimer’s Disease Neuroimaging Initiative (see note)

**Harmonization of methods**: Alzheimer’s Disease Metabolomics Consortium (see note)

**Technical, bibliographic research and/or material support**: Louie

**Critical revision of the manuscript for important intellectual content**: Saykin, Kaddurah-Daouk

**Obtained funding**: Kaddurah-Daouk

**Supervision**: Saykin, Kastenmüller, Kaddurah-Daouk

**The Alzheimer’s Disease Neuroimaging Initiative (ADNI)**: Data used in the preparation of this article were obtained from the ADNI database (http://adni.loni.usc.edu). As such, the investigators within the ADNI contributed to the design and implementation of ADNI and/or provided data but did not participate in analysis or writing of this report. A complete listing of ADNI investigators can be found at: http://adni.loni.usc.edu/wp-content/uploads/how_to_apply/ADNI_Acknowledgement_List.pdf.

## Funding/Support

Funding for ADMC (Alzheimer’s Disease Metabolomics Consortium, led by Dr R.K.-D. at Duke University) was provided by the National Institute on Aging grant 1U01AG061359-01and R01AG046171, a component of the Accelerating Medicines Partnership for AD (AMP-AD) Target Discovery and Preclinical Validation Project (https://www.nia.nih.gov/research/dn/amp-ad-target-discovery-and-preclinical-validation-project) and the National Institute on Aging grant RF1 AG0151550, a component of the M^2^OVE-AD Consortium (Molecular Mechanisms of the Vascular Etiology of AD – Consortium https://www.nia.nih.gov/news/decoding-molecular-ties-between-vascular-disease-and-alzheimers).

Data collection and sharing for this project was funded by the Alzheimer’s Disease Neuroimaging Initiative (ADNI) (National Institutes of Health Grant U01 AG024904) and DOD ADNI (Department of Defense award number W81XWH-12-2-0012). ADNI is funded by the National Institute on Aging, the National Institute of Biomedical Imaging and Bioengineering, and through generous contributions from the following: AbbVie, Alzheimer’s Association; Alzheimer’s Drug Discovery Foundation; Araclon Biotech; BioClinica, Inc.; Biogen; Bristol-Myers Squibb Company; CereSpir, Inc.; Eisai Inc.; Elan Pharmaceuticals, Inc.; Eli Lilly and Company; EuroImmun; F. Hoffmann-La Roche Ltd and its affiliated company Genentech, Inc.; Fujirebio; GE Healthcare; IXICO Ltd.; Janssen Alzheimer Immunotherapy Research & Development, LLC.; Johnson & Johnson Pharmaceutical Research & Development LLC.; Lumosity; Lundbeck; Merck & Co., Inc.; Meso Scale Diagnostics, LLC.; NeuroRx Research; Neurotrack Technologies; Novartis Pharmaceuticals Corporation; Pfizer Inc.; Piramal Imaging; Servier; Takeda Pharmaceutical Company; and Transition Therapeutics. The Canadian Institutes of Health Research is providing funds to support ADNI clinical sites in Canada. Private sector contributions are facilitated by the Foundation for the National Institutes of Health (www.fnih.org). The grantee organization is the Northern California Institute for Research and Education, and the study is coordinated by the Alzheimer’s Disease Cooperative Study at the University of California, San Diego. ADNI data are disseminated by the Laboratory for Neuro Imaging at the University of Southern California.

The work of various Consortium Investigators are also supported by various NIA grants [U01AG024904-09S4, P50NS053488, R01AG19771, P30AG10133, P30AG10124, K01AG049050], the National Library of Medicine [R01LM011360, R00LM011384], and the National Institute of Biomedical Imaging and Bioengineering [R01EB022574]. Additional support came from Helmholtz Zentrum, the Alzheimer’s Association, the Indiana Clinical and Translational Science Institute, and the Indiana University-IU Health Strategic Neuroscience Research Initiative.

**Role of the Funder/Sponsor**:[Funders listed above] had no role in the design and conduct of the study; collection, management, analysis, and interpretation of the data; preparation, review, or approval of the manuscript; and decision to submit the manuscript for publication.

**Additional Contributions**: The authors are grateful to Lisa Howerton for administrative support and the numerous ADNI study volunteers and their families.

