## Supplemental Table 1 for "Serum Metabolites Associated with Brain Amyloid Beta Deposition, Cognitive Dysfunction, and Alzheimer’s Disease Progression"

### Supplementary Table 1 – Demographics of ADNI participants stratified by baseline diagnosis

| | CN (N=370) | SMC (N=95) | EMCI (N=271) | LMCI (N=491) | AD (N=304) |
|---|---|---|---|---|---|
| Age | 74.6 (5.8) | 72.3 (5.7) | 71.3 (7.6) | 74.0 (7.6) | 74.8 (7.8) |
| Sex: (Male/Female) | 181/189 | 40/55 | 149/122 | 299/192 | 166/138 |
| Education, years | 16.3 (2.8) | 16.7 (2.6) | 16.0 (2.7) | 15.8 (2.9) | 15.2 (3.0) |
| BMI (Kg/M$^2$) | 27.0 (4.5) | 28.4 (6.2) | 28.0 (5.4) | 26.4 (4.3) | 25.9 (4.7) |
| *APOE* ε4 (0/1/2) | 266/95/9 | 66/28/1 | 156/96/19 | 224/201/66 | 106/139/59 |
| Memory composite | 1.03 (0.57) | 1.08 (0.54) | 0.58 (0.60) | -0.08 (0.62) | -0.85 (0.54) |
| Executive function composite | 0.78 (0.79) | 0.79 (0.85) | 0.50 (0.85) | 0.01 (0.86) | -0.90 (0.93) |
| MMSE | 29.1 (1.1) | 29.0 (1.2) | 28.3 (1.60) | 27.1 (1.83) | 23.3 (2.0) |
| C3 | -0.02 (1.01) | -0.06 (0.73) | -0.15 (0.91) | -0.04 (0.90) | -0.23 (0.93) |
| Kynurenine | -0.08 (0.98) | -0.31 (0.87) | -0.32 (0.88) | -0.12 (0.96) | -0.24 (1.04) |
| LysoPC a C18:2 | 0.17 (0.96) | -0.11 (0.89) | 0.04 (0.92) | 0.37 (0.97) | 0.27 (0.92) |
| PC aa C42:0 | 0.11 (0.98) | 0.01 (1.01) | 0.01 (0.90) | 0.26 (0.95) | 0.24 (0.91) |
| PC ae C42:3 | 0.44 (0.91) | 0.17 (1.00) | 0.27 (0.95) | 0.49 (0.91) | 0.40 (0.92) |
| PC ae C44:3 | 0.34 (0.92) | 0.28 (0.97) | 0.07 (0.91) | 0.45 (0.93) | 0.41 (0.94) |
| PC ae C44:4 | 0.28 (0.95) | 0.12 (0.96) | 0.17 (0.99) | 0.46 (0.93) | 0.45 (0.91) |
| PC ae C44:5 | 0.15 (1.01) | 0.02 (0.94) | 0.09 (1.01) | 0.30 (0.94) | 0.32 (0.90) |
| PC ae C44:6 | 0.13 (1.00) | -0.03 (0.96) | 0.06 (0.94) | 0.34 (0.94) | 0.31 (0.92) |

*Data are reported as mean (SD) unless otherwise indicated.
Abbreviations: AD: Alzheimer's disease; BMI: Body mass index; CN: Cognitively normal older adults; EMCI: Early mild cognitive impairment; LMCI: Late mild cognitive impairment; SMC: subjective memory complaint; ADAS-Cog 13: modified 13-item Alzheimer's Disease Assessment Scale, cognitive subscale; CDR-SB: Clinical Dementia Rating-Sum of Boxes; MMSE: Mini-Mental State Examination;
